# Dorsoventral limb patterning in paired appendages emerged via regulatory repurposing of an ancestral posterior fin module

**DOI:** 10.1101/2025.04.16.648507

**Authors:** Sofía Zdral, Simone Giulio Bordignon, Axel Meyer, Marian A. Ros, Joost M. Woltering

## Abstract

Limbs exhibit adaptive differentiation along their dorsoventral (DV) axis, determined by the dorsal expression of the LIM homeobox gene *Lmx1b*. The paired appendages (i.e. the pectoral and pelvic fins from which limbs evolved) arose in an early jawless ancestor via co-option of a midline-fin genetic program including modules for anteroposterior (AP) and proximodistal (PD) patterning. Unlike the AP and PD axes, median fins lack an unambiguous DV axis, leaving the origin of this DV pattern in paired appendages unresolved. Here, we describe *Lmx1b* expression in the posterior midline fins of cichlids, sturgeons and catsharks, revealing an ancestral role for this gene predating the origin of paired appendages. In median fins, *Lmx1b* activation depends on *shh* from the ZPA, whereas in paired fins it relies on ectodermal *wnt* signalling, indicating the evolution of novel regulatory inputs for dorsal patterning. We observe *ephA4b*, a putative *Lmx1b* target, is co-expressed with *Lmx1b* in dorsal pectoral and posterior midline fins and downregulated alongside *Lmx1b*, suggesting a role in both fin types related to axon guidance. We propose that novel regulation drove the repurposing of *Lmx1b* from posterior to dorsal fin determinant, with co-option of conserved downstream targets. Altogether, our findings demonstrate that the DV axis of paired appendages represents an evolutionary innovation arising from the integration of ancestral midline fin and flank determinants with novel regulatory inputs.

## Introduction

The paired appendages, i.e. forelimbs and hindlimbs, and their evolutionary precursors the pectoral and pelvic fins, evolved in a jawless ancestor around 500 million years ago (MYA) (Coates 1994; Bayramov, et al. 2024). Prevailing theories explain this event by co-option of the midline fins to the flanks (Freitas, et al. 2006; Onimaru, et al. 2011; Tanaka and Onimaru 2012; Gai, et al. 2022; Tzung, et al. 2023; Bayramov, et al. 2024), potentially facilitated by the gill arches (Gillis, et al. 2009; Diogo 2020; Sleight and Gillis 2020; Brazeau, et al. 2023). This step marks a pivotal transition in vertebrate evolution without which weight-bearing terrestrial locomotion would have been impossible to achieve. Coherent functioning of the paired fins and limbs relies on the correct integration of developmental processes along their three main axes (**Fig. 1**). The genetic modules for the anteroposterior (AP) axis (running perpendicular to the digits or fin rays (Woltering, et al. 2020)) and proximodistal (PD) axis (from body to appendage tip (Woltering and Duboule 2010)) are shared between median and paired fins and reflect their common origin in the midline (Freitas, et al. 2006; Letelier, et al. 2018; Höch, et al. 2021; Hawkins, et al. 2022). While these two axes can thus be considered as homologous between the two types of appendages, this is less clear for the third axis - the dorsoventral (DV) axis (Castilla-Ibeas, et al. 2024). This latter polarity is crucial for locomotion as it differentiates a specialized substrate interacting-earth facing-surface together with counteracting extensor and flexor muscle groups (Castilla-Ibeas, et al. 2024) (**Fig. 1**), which in land animals are required to lift the body off the ground. Such adaptive differentiation of the limbs is evident throughout the tetrapod radiation (land vertebrates) and in mammals this is further manifested by the presence of footpads and nails, which originate from the respective ventral and dorsal sides of hand and feet (Castilla-Ibeas, et al. 2024). Evidence of DV specialization predates the tetrapod lineage as tetrapodomorph fish exhibit specialized ventral hemi-rays in their pectoral and pelvic fins (Stewart, et al. 2020), and a pattern of opposing abductor and adductor muscles with dedicated motor neuron signature is present in fins of fish (Uemura, et al. 2005; Ziermann, et al. 2017; Jung, et al. 2018; Siomava and Diogo 2018; Siomava, et al. 2018; Hoffmann, et al. 2019).

**Fig. 1.**
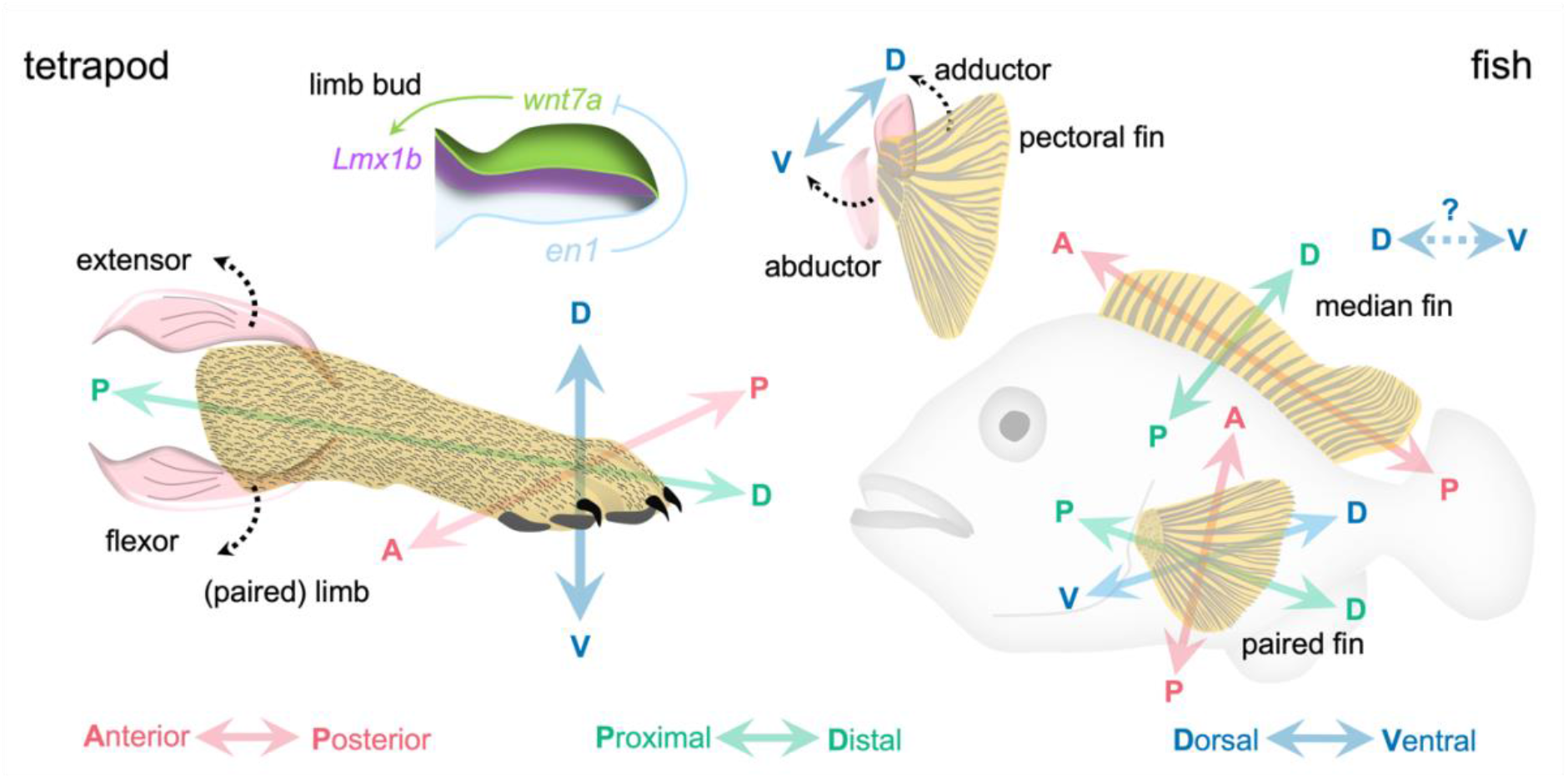
Axial polarities in fins and limbs. Three axes are recognized in paired fins whereby the dorsoventral axis (DV, blue), in mammals defines a ventral substrate interacting surface and nails arising from the dorsal surface, as well as opposing extensor and flexor muscles. Dorsal identity is specified by expression of the Lmx1b master control gene during development. Similar dorsoventral organization is present in fish paired fins. Median fins, in contrast. have homologous anteroposterior (AP, red) and proximodistal (PD, green) polarity but no unambiguous dorsoventral axis. Therefore, the relationship between the Lmx1b patterning module in paired and median fins as well as the evolutionary origin of the DV axis in paired appendages remains unknown.

DV differentiation is driven by the exclusively dorsal expression of the *Lmx1b* LIM homeodomain gene, a master regulator of dorsal identity (Chen, et al. 1998; Chen and Johnson 2002; Castilla-Ibeas, et al. 2024). *Lmx1b* loss of function in mouse results in double ventral limbs with footpads on both sides and absence of nails (Castilla-Ibeas, et al. 2024). In humans, mutations in this gene are related to Nail-Patella Syndrome (OMIM #161200) with patterning abnormalities similar to those found in the *Lmx1b*-null mice (Chen, et al. 1998; Chen and Johnson 2002; Castilla-Ibeas, et al. 2024). In addition, *Lmx1b* is required for the correct innervation of dorsal extensor muscles via regulation of downstream ephrin receptors (Kania, et al. 2000; Jung, et al. 2018). The upstream network for dorsal *Lmx1b* expression relies on ectodermal activity of *wnt7a*. Ventrally, this pathway is repressed by *engrailed1* (*en1*) (**Fig. 1**) (Parr and McMahon 1995; Riddle, et al. 1995; Cygan, et al. 1997; Chen and Johnson 2002).

Mutations in *en1* or *wnt7a* result in (distally) bi-dorsal and bi-ventral limbs, respectively, consistent with their role in establishing the DV axis (Parr and McMahon 1995; Riddle, et al. 1995; Yang and Niswander 1995; Loomis, et al. 1996; Cygan, et al. 1997; Loomis, et al. 1998; Chen and Johnson 2002). The evolutionary conservation of this DV patterning network is inferred from the dorsal *Lmx1b* expression in zebrafish and skate pectoral fins (Uemura, et al. 2005; Jung, et al. 2018). This dorsal *Lmx1b* fin domain expectedly contributes to the general DV organization, including tetrapod-like motor neuron innervation of the pectoral adductor and abductor muscles, resembling that of the limb‘s extensors and flexors (Uemura, et al. 2005; Ziermann, et al. 2017; Jung, et al. 2018; Siomava and Diogo 2018; Siomava, et al. 2018; Hoffmann, et al. 2019). Whereas the AP and DV patterning of the paired fins and limbs can be traced directly to an ancestral midline fin pattern, the origins of the DV axis remain elusive. The orthogonality of the AP, PD and DV axes indicates that the DV axis in paired fins should correspond to the midline fin‘s left-right division (**Fig. 1**). Since midline fins follow the symmetry axis of the bilaterian body plan, it is uncertain if they possess any lateral polarity. When the paired fins arose, they however became aligned with the DV plane of the trunk and could implement an asymmetry absent from their midline precursors. The adaptive value of the specialized substrate/gravity and water surface facing sides of the paired fins is obvious indeed, unlike that of any left-right differentiation of a midline fin. The AP and PD axes of paired fins thus have clear evolutionary antecedents in the midline, and their orientation towards both the body and external environment mirrors this ancestral condition. However, the DV axis presents a more complex case, with its origins and relationship to pre-existing midline fin modules remaining uncertain. It has been established that the ventral aspect of fins and limbs likely derives from an ancestral DV division in the trunk that predates the origin of paired fins (Tanaka, et al. 2002; Matsuura, et al. 2008; Tanaka 2013). However, how *Lmx1b* and its upstream regulatory pathways co-ordinately arose as dorsal determinants, and whether these bear any relationship to ancestral trunk or midline fin modules remains unresolved.

*Lmx1b* is a highly pleiotropic gene with roles not only in limb DV patterning but also in the development of the central nervous system (CNS) and kidneys (Chen, et al. 1998; Haldin, et al. 2008) from which tissues it may have been co-opted after paired appendages arose. Alternatively, *Lmx1b* has lesser known functions in the limb which extend beyond an exclusive role in DV patterning (Chen and Johnson 2002). For instance, ulnar dysplasia is consistently observed in *Lmx1b* null mouse suggesting an additional role in AP patterning (Chen, et al. 1998; Chen and Johnson 2002). Furthermore, it is known that *Lmx1b* acts redundantly with two other LIM domain proteins, *Lhx2* and *Lhx9*, to integrate signals along the AP, PD and DV axes of the limbs (Tzchori, et al. 2009). This suggests that *Lmx1b* could have had an ancestral role in the median fins unrelated to DV patterning, which was expanded upon co-option to the flanks.

Understanding the evolutionary trajectory of acquisition of *Lmx1b* expression in paired appendages, and its relationship to midline fins, is key to understanding how coherent fins and limbs arose. To this end we comparatively investigated *Lmx1b* midline expression and regulation in a suite of fishes selected for their phylogenetic position. We chose to investigate expression in a teleost species (the group of fish considered to be most “modern” and comprising 99% of current fish diversity), namely the cichlid *Astatotilapia burtoni*. This species has an advantage over other models that it is “directly developing” (Woltering, et al. 2018), and all fins, including midline fins, differentiate early during development (opposed to zebrafish (Parichy, et al. 2009)). This is for instance demonstrated by the activation of the posterior *shh*-*gremlin* patterning module in dorsal and anal fins already within the first week of development (Höch, et al. 2021), and facilitates a comparison of paired and midline appendages at similar stages. We further selected the sturgeon (*Acipenser baerii*) (Fopp-Bayat, et al. 2022), which is an early branching ray finned fish that diverged from the lineage of the teleosts around 345 MYA (Du, et al. 2020). We also investigated *Lmx1b* expression in the small spotted catshark (*Scyliorhinus canicula)* (Ballard, et al. 1993), which is part of the *Chondrichthyes* (cartilaginous fishes) rather than the *Osteichthyes* (bony fishes) from which it split around 450 MYA (Hara, et al. 2018). Shared characters between these lineages would indicate presence in their last common ancestor close to the origin of the *Gnathostomes* (jawed vertebrates) and the origin of the paired appendages.

## Results

### *Lmx1b* is expressed in posterior median fins and unrelated to *en1* or *wnt7a*

We investigated expression of *Lmx1b* in *A. burtoni* from 3-8 days post fertilization (dpf) (**Fig. 2a, Fig.S1, Fig.S2**), covering a time window during which the *Anlagen* for both paired and midline fins are established (Woltering, et al. 2018). As a result of the teleost specific genome duplication (3R) two genomic *Lmx1b* gene copies are present – *Lmx1ba* and *Lmx1bb. Lmx1ba* only appears expressed in the anterior CNS and forming gut but is absent from the appendages (**Fig.S3**). *Lmx1bb* is however expressed in both paired and midline fins, indicating that the *Lmx1ba* paralog lost its role in fin patterning due to sub-functionalisation. In the emerging pectoral and pelvic fins, *Lmx1bb* is expressed in the expected dorsal mesenchymal domain as soon as the fin bud emerges (**Fig. 2a, Fig.S1, Fig.S2**). In the midline fins we observe previously unreported expression in both the dorsal and anal fins in a posterior domain coinciding with the Zone of Polarizing Activity (ZPA) as evidenced by the expression of *shh* (Höch, et al. 2021). No expression was detected in the caudal fin, which also lacks a distinguishable ZPA. In the emerging pectoral and pelvic fin buds, *wnt7aa* and *en1b* show superficial, dorsal and ventral expression respectively (**Fig. 2a, Fig.S1, Fig.S2**), consistent with their known roles in restricting *Lmx1b* to the dorsal side. At none of the stages investigated (3-8 dpf) do we observe *en1b* expression in the midline fins (**Fig.2a, Fig.S1)** except for a weak domain in the posterior fin folds at 8dpf (**Fig.S2)**. In the median fins, *wnt7aa* is expressed in a distal ectodermal domain that initially extends along the AP axis and becomes subsequently restricted to the prospective adult dorsal midline fin (as contrasting with the fin fold) (**Fig.2a, Fig.S1**). Dorsal views show the superficial (ectodermal) expression of *wnt7aa*, which contrasts with the expression of *Lmx1bb* more towards the centre of the midline (**Fig.2a**).

**Fig. 2.**
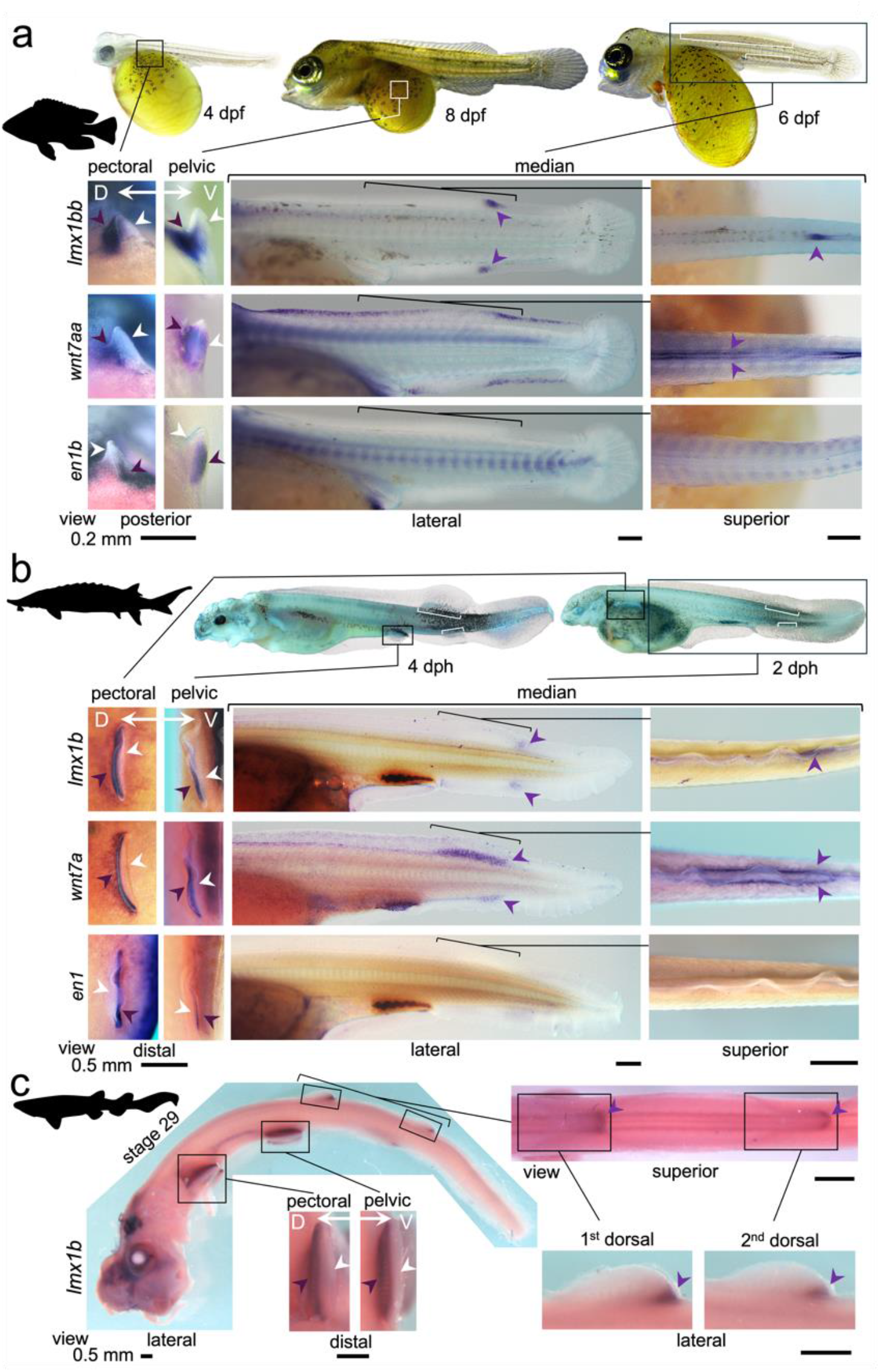
Expression of Lmx1b, wnt7a and en1. **a**) Expression in A. burtoni in 4, 6 and 8 dpf embryos. **b**) Expression in A. baerii in 2 and 4 dph embryos. **c**) Expression in castshark st.29. Because of heterochronic development of the pectoral, pelvic and median appendages these are shown for different developmental stages in A.burtoni and A. baerii. Lilac arrowheads indicate sites of notable expression while white arrowheads indicate absence of expression. White brackets in overviews of 6dpf A.burtoni (**a**) and 2/4dph A. baerii (**b**) indicate the extent of the forming adult dorsal and anal fins, distinguishing those from the fin folds. Dpf: days post fertilization; dph: days post hatch; st: stage.

The expression of *Lmx1b* in the posterior dorsal and anal fins and its association with the ZPA suggests a function that predates the co-option of the median fin module to the flank. To confirm that this expression domain is ancestral to gnathostomes - as opposed to having a derived apomorphic origin in modern teleost fish - we further investigated this in a comparative context using sturgeon (*A. baerii*) and catshark (*S. canicula*) embryos. In the pectoral and pelvic fins of 2/4 days post hatching (dph) sturgeon we observed the expected expression domains along the DV axis whereby *Lmx1b* and *wnt7a* are expressed dorsally, while *en1* is ventrally restricted (**Fig.2b**). In the dorsal and anal fin *Anlagen* of 2dph embryos we observe a very similar pattern compared to that in *A. burtoni*, with absence of *en1*, expression of *wnt7a* corresponding to the entire adult fin forming domains (see white brackets in embryo overviews (**Fig.2b)** to distinguish these from anterior and posterior fin folds), and a posteriorly restricted domain of *Lmx1b*. In the *S. canicula* we investigated expression of *Lmx1b*, which in pectoral and pelvic fins of Ballard stage 28 and 29 (Ballard, et al. 1993) is expressed in the dorsal half of the fin mesenchyme (**Fig.2c, Fig.S4**) as well as at the posterior border of dorsal and anal fins in a domain reminiscent to that present in sturgeon and *A. burtoni*.

Altogether, this confirms that expression of *Lmx1b* in the dorsal paired appendages is ancestral for jawed vertebrates (Uemura, et al. 2005; Jung, et al. 2018). Moreover, the identification of a previously undescribed domain of *Lmx1b* expression in the posterior border of midline fins, shared among the three investigated gnathostome lineages, strongly indicates a midline fin role in jawed vertebrates, that was present prior to the emergence of the flank appendages.

### Midline fin *Lmx1bb* relies on *shh* but not on *wnt* signalling

In paired appendages, *Lmx1b* is dorsally activated by dorsal ectodermal *wnt7a* expression (Parr and McMahon 1995; Riddle, et al. 1995; Cygan, et al. 1997; Chen and Johnson 2002), the expression of which we show to similarly correlate with *Lmx1b* in pectoral and pelvic fins. In the median fins however, the much more extended AP *wnt7a* domain as compared to *Lmx1b* suggests that a different relationship between *wnt* signalling and *Lmx1b* expression exists in the midline. This suggest that *Lmx1b* may rather be regulated by a different, exclusively posterior signal, such as *shh* derived from the ZPA (Höch, et al. 2021). We therefore investigated the roles of *wnt* and *shh* signalling in activating *Lmx1bb* in the pectoral, pelvic and midline fin domain in *A. burtoni* by chemical modulation of these pathways. *Shh* was inhibited using cyclopamine or activated using SAG (Höch, et al. 2021) and *wnt* signalling was inhibited using IWR-1-endo (Kimura, et al. 2022). Treatments were performed for 2 days during the initial stages of fin formation, which correspond to 2-4 dpf for pectoral fins, 6-8 dpf for pelvic fins and 4-6 dpf for midline (dorsal and anal) fins (Woltering, et al. 2018). As BMP signalling is known to play an important role in the establishment of the AP division in median fins, we also investigated this pathway using the BMP inhibitor DMH1 as well as fish mutant for the secreted BMP antagonist *gremlin1b* (Höch et al. 2021).

In pectoral and pelvic fins no changes in *Lmx1bb* expression were observed after manipulation of the *shh* pathway (**Fig. 3a, b**) while suppression of *wnt* signalling resulted in downregulation of *Lmx1bb* expression (**Fig. 3a, b**), similar as observed for downregulation of *wnt* signalling in mouse limbs (Parr and McMahon 1995; Riddle, et al. 1995). In dorsal and anal fins, the opposite effects were observed (**Fig. 3c**): suppression of *wnt* signalling did not alter *Lmx1bb* expression, while inhibition of *shh* signalling caused strong downregulation of *Lmx1bb* expression at the position of the ZPA. Intriguingly, activation of the *shh* pathway only resulted in a very slight upregulation of *Lmx1bb* expression in the most anterior part of the dorsal fin (**Fig.3c**). This suggests that *shh* signalling, although required, is only one part of a more complex upstream activating module. Because BMP signalling is known to play an important role in the patterning of dorsal and anal fins, we investigated potential crosstalk with *shh* signalling in the regulation of *Lmx1bb*. Inhibition of BMP signalling using DMH1 did not result in differences in expression, while in mutants for the secreted BMP antagonist *gremlin1b* a modest anterior expansion of the *Lmx1bb* domain was observed (**Fig.3c**), suggesting a role for BMP in anteriorly restricting the *Lmx1bb* domain. Interestingly, it is known that *gremlin1b* is highly upregulated by *shh* signalling in median fins (Höch, et al. 2021), suggesting a possible explanation for the inability of SAG to upregulate *Lmx1bb* expression in wildtype (WT) genetic background. We therefore stimulated *shh* signalling in *gremlin1b* knockout mutant embryos. Indeed, under these conditions *Lmx1bb* expression became expanded throughout the AP extent of the dorsal and anal fins (**Fig.3c**), indicating that the posterior midline fin domain depends on complex regulatory interactions between BMP and *shh* pathways. Hereby, *shh* acts as the main activating agent of *Lmx1bb* in the posterior most fin domain, whose anterior boundary seems to be controlled by the expression of *gremlin1b*.

**Fig. 3.**
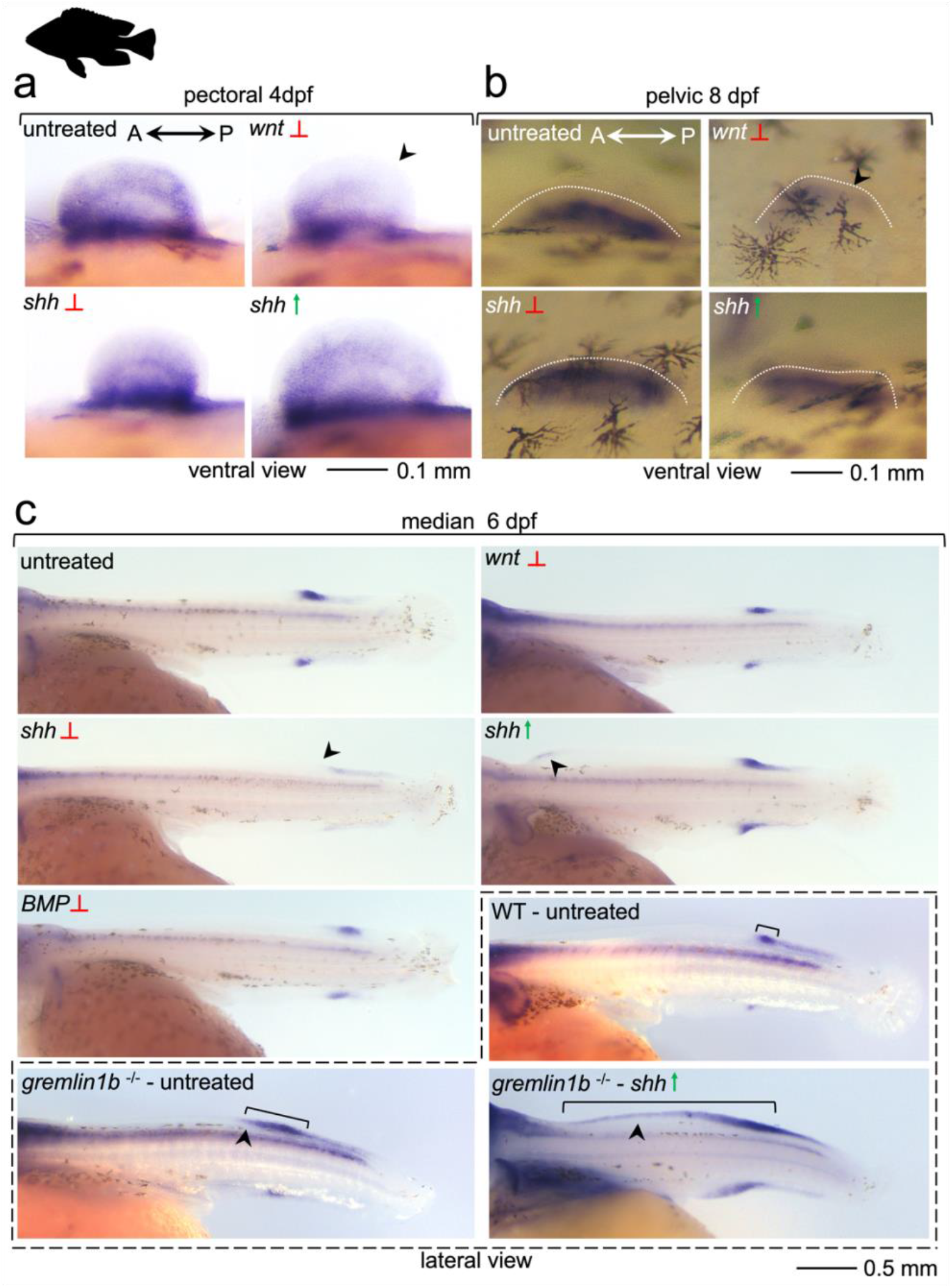
Differential regulation of Lmx1bb in A. burtoni paired and median fins. Pectoral (**a**), pelvic (**b**) or midline (dorsal and anal) (**c**) fins were treated using modulators of the shh or wnt signal transduction pathways. Green arrows indicate agonist treatment, while red braking signs indicated inhibitory treatments. Black arrowheads are used to indicate notable changes in Lmx1bb expression referred to in the main text. Downregulation of Lmx1bb can be observed with inhibition of wnt signalling in pectoral and pelvic fins; and using shh inhibition in median fins. In median fins the role BMP signalling was investigated using gremlin1b knockout mutants, which show a modest anterior expansion of expression and a strong anterior expansion upon treatment with shh agonist (compare with brood control within dashed lines).

Most importantly, we experimentally demonstrate that different signals are responsible for the activation of *Lmx1bb* in paired and midline appendages; canonical *wnt* signalling drives expression in the paired fins as in tetrapod limbs, whereas the posterior domain in dorsal and anal fins is dependent on *shh* secreted from the ZPA. Altogether, this suggests that regulatory changes in *Lmx1bb* accompanied its emergence as dorsal determinant during the evolution of paired appendages.

### Evolutionary history of a LIM module during the emergence of paired limbs

In addition to its role as a DV selector gene, *Lmx1b* is part of a transcriptional module of LIM-HD factors that functions to integrate signals for the AP and PD patterning of the limbs (Tzchori, et al. 2009).

Within this module, it works redundantly with the related LIM-HD genes *Lhx9* and *Lhx2* for whose absence it can compensate dorsally. Such role of *Lmx1b* in the patterning of the PD and AP axes as part of a LIM-HD transcriptional module could represent an ancestral role in median fins, which became modified to DV regulator after emergence of new regulation in the paired appendages. We investigated how the fin expression of *Lmx1b* compares to that of *Lhx2* and *Lhx9* and whether similarity of expression in the midline fins would be supportive of such scenario. In the pectoral fins of *A. burtoni* and sturgeon *Lhx2(b)* and *Lhx9* show similar expression domains, as reported for mouse limbs (**Fig. 4a)** (Tzchori, et al. 2009), with more or less ubiquitous distribution along the AP axis and a somewhat stronger expression towards the distal margin. In the dorsal and anal fins, these genes are expressed in a similar domain along the entire extent of the AP axis, with *Lhx9* showing slightly weaker expression posteriorly, while *Lhx2(b)* shows a stronger signal distally. In sturgeon, *Lhx2* appears excluded from the posterior pectoral fin and in *A. burtoni Lhx9* is expressed in lower intensity in the posterior dorsal fin, showing some species specific differences as also reported for chicken and mouse limbs, which is consistent with their strong redundancy (Tzchori, et al. 2009). Furthermore, and in contrast to *Lmx1b*, these two genes are also expressed in the caudal fin (**Fig.4a**). Altogether, and unlike is the case for *Lmx1b, Lhx9* and *Lhx2* show strong similarity in their spatial expression in the paired and midline fins and are expressed in a domain consistent with their described role as integrating signals for AP and PD patterning in the limbs. This similarity between paired and midline fins suggests a conserved role across these fin types that was present before the appearance of the paired fins and was part of an ancestral module co-opted to the flank. However, the strong differences in midline fin expression between *Lhx2*/*Lhx9* and *Lmx1b* suggests, that the redundant incorporation of *Lmx1b* into a “LIM module” is likely not ancestral but only occurred together with the anterior expansion of its expression domain in the paired appendages and would therefore not represent an ancestral midline function.

**Fig. 4.**
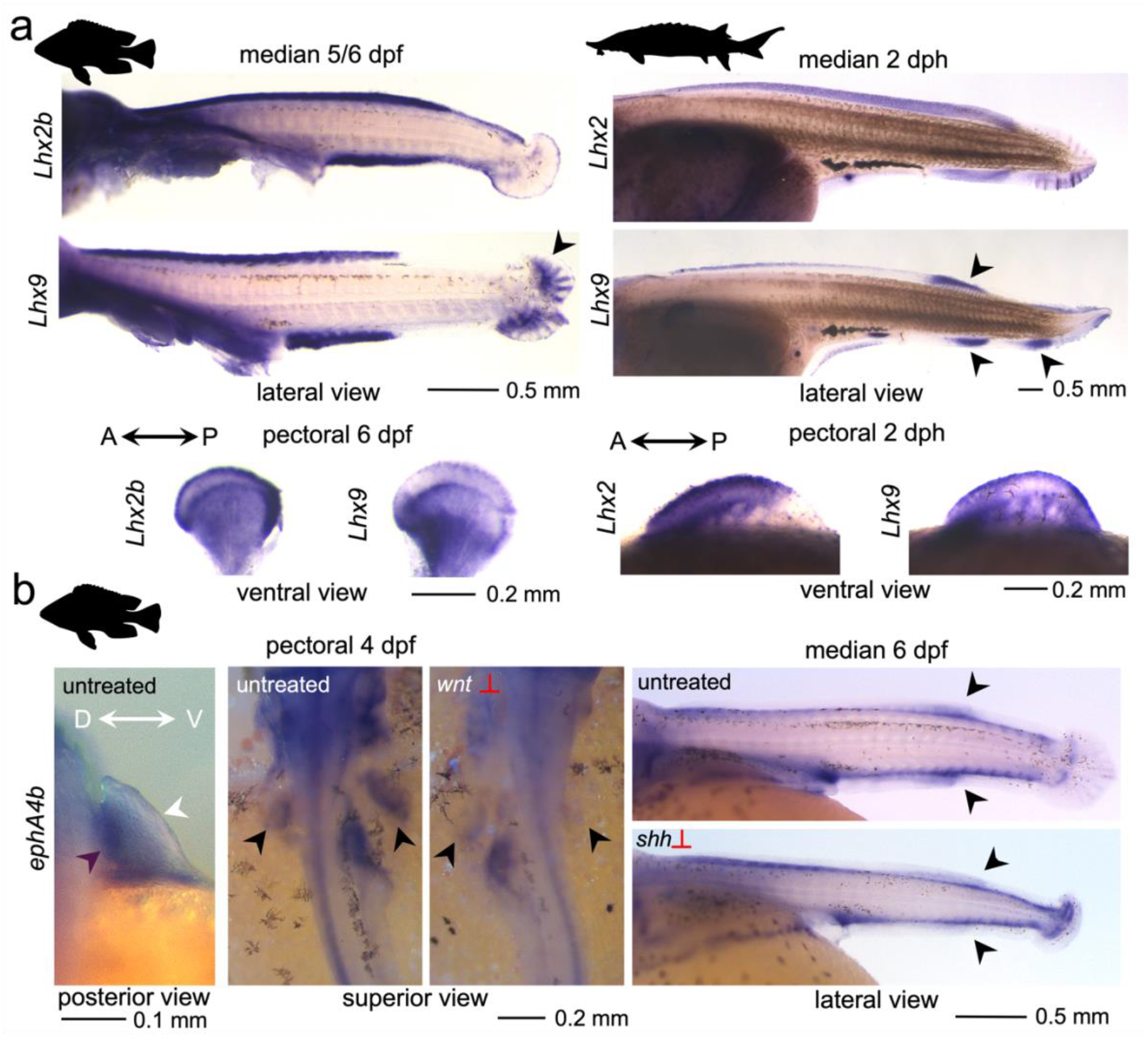
Lhx2/Lhx9 and ephA4b receptor expression. **a**) Expression of Lhx2(b) and Lhx9 in A. burtoni (left) and A. baerii (right) in different fins (median and pectoral) and at different stages. **b**) Fin expression of ephA4b in A. burtoni. At 4dpf ephA4b is expressed in the dorsal part of the pectoral fins and downregulated upon inhibition of wnt signalling. Downregulation is additionally observed in neural tissues, but not in the forming gut. In median fins inhibition of shh signalling leads to downregulation of ephA4b. This corresponds to downregulation of Lmx1bb at these sites (**Fig.S5**).

### Conserved overlap of *ephA4b* and *Lmx1bb* expression in paired and midline fins

A major role of the DV pattern in limbs is to coordinate motor neuron innervation in the dorsal and ventral halves of the limbs to specify extensor and flexor type muscles. *Lmx1b* has been shown to regulate this process by controlling ephrin receptor *ephA4* expression dorsally, the downregulation of which in *Lmx1b* mutant mice results in incorrect innervation and muscle function (Kania, et al. 2000; Kania and Jessell 2003). A similar pattern of dorsal ephrin receptor expression and motor neuron migration in fish is related to dorsal *Lmx1b* expression (Jung, et al. 2018). Little is as yet understood about the homology between median and paired fin muscles (see discussion in (Siomava and Diogo 2018)) and no direct homologies are established between the two fin types. Yet, the coordination of motor neuron innervation of posterior specific median fin muscles, such as the *retractor dorsalis*/*retractor analis* (Siomava, et al. 2020), would provide a plausible ancestral function for the posterior *Lmx1b* midline fin expression domain. We investigated this hypothesis by looking at the overlap of ephrin receptor expression in dorsal paired and posterior midline fins. One of the ephrin receptors dorsally downregulated in *Lmx1b* mouse mutant limbs (Kania, et al. 2000) and dorsally expressed in paired fish fins (Jung, et al. 2018) is *ephA4*. Indeed, in *A. burtoni ephA4b* is dorsally expressed in pectoral fins (**Fig. 4b**). In dorsal and anal fins, it is expressed in a posterior domain correlating with the expression domain of *Lmx1bb*, suggesting it as a conserved target. This was further investigated using the conditions shown to downregulate *Lmx1bb* expression. In line with the downregulation of *Lmx1bb* in *wnt* inhibited and *shh* inhibited conditions in pectoral and median fins respectively (**Fig. 3, Fig. S5**), a concomitant downregulation of *ephA4b* is observed, further strengthening the connection between *Lmx1bb* and *ephA4b* expression domains in the fins. Altogether, these observations would provide support for an ancestral midline function of *Lmx1bb* expression in coordinating differential innervation of anterior versus posterior domains of the midline fins.

## Discussion

### The limbs DV axis as evolutionary novelty

Unlike the origin of the proximodistal and anteroposterior axes, the dorsoventral axis of the paired appendages cannot be directly interpreted as resulting from the co-option of one of the ancestral midline fin axes. Here, we show that the primary determinant of the paired appendage DV axis, *Lmx1b*, is expressed in a conserved posterior midline fin domain that predates the origin of the paired appendages. Analysis of upstream signalling identifies *shh* as the primary activator in the midline fin. This contrasts with the paired appendages where the dorsal domain of *Lmx1b* is under control of canonical *wnt* signalling. In combination with the strong differences in AP distribution of *Lmx1b* transcription in paired versus dorsal/anal fins, this suggests that the ancestral midline fin domain cannot be directly homologized with its dorsal domain in the paired fins. Altogether these findings indicate that the DV axis arose as a novelty in paired appendages via the evolution of new dorsal regulatory inputs such as potentially the *Lmx1b* limb specific *LARM* enhancers (Haro, et al. 2021). Given the broad distribution of *wnt* signalling in the median fins – and embryo as a whole-, such enhancers must show strong context specificity for the lateral plate mesoderm (LPM) and therefore likely involve multiple other activating and restricting upstream factors.

### Integrating ancestral gene functions with novel regulations

At the same time, the conserved posterior midline fin domain of *Lmx1b* suggests an ancestral role that predates the evolution of paired appendages (**Fig. 5**). The posterior nature of the expression in the midline fins aligns intriguingly with the observed posterior skeletal limb defects in *Lmx1b* mutant mice (Chen, et al. 1998; Chen and Johnson 2002) and in NPS patients (Dreyer, et al. 2000; Sweeney, et al. 2003) which therefore may be linked to its ancestral function. However, given the strong divergence in spatial expression in the median fins between *Lmx1b* and L*hx2*/*Lhx9*, such ancestral role of *Lmx1b* seems likely unrelated to its current integration into a module of LIM domain genes acting in AP and DV patterning (Tzchori, et al. 2009). Concerning the acquisitions of its function as dorsal master regulator, it is plausible that ancestral downstream targets of *Lmx1b* were co-opted dorsally in the paired appendages and provided the selective basis for its transition from a posterior to a dorsal determinant. A clear example of this is the activation of ephrin receptor signalling for motor neuron guidance, as we find the expression of *ephA4b* to widely correlate with the expression of *Lmx1bb*. Interestingly, this is not restricted to the fins but includes neural tissue, where *Lmx1bb* and *ephA4b* are similarly downregulated by *wnt* inhibition. Furthermore, we observe *ephA4b* expression in the gut to overlap with that of *Lmx1ba* (**Fig.4b**/**Fig.S3**). Under such scenario, the evolution of novel *wnt* responsive regulation for dorsal *Lmx1b* expression in the paired fins would directly provide a substrate for adaptive diversification into extensor and flexor type muscles via the activation of a conserved downstream target. The ventral *en1* expression in paired appendages appears co-opted from an ancestral division between ventral and dorsal parts of the LPM on which border the paired fins have emerged (Tanaka, et al. 2002; Tanaka and Onimaru 2012; Tanaka 2013). Here, we show that superficial expression of *wnt7a* is present in forming *A. burtoni* and *A. baerii* midline fins and therefore likely was part of the original median fin module that became co-opted to the flanks (**Fig.5**). It is however unclear how the repressive interactions between *en1* and *wnt7a* arose to restrict the latter to a dorsal domain. It is possible this repressive interaction already plays an ancestral role in a different tissue context or, alternatively, that it arose after emergence of the paired appendages. Dorsal limbs have been induced in chicken dorsal midline (Yonei-Tamura, et al. 1999) where symmetrical *Lmx1b* expression is induced by a dorsal *wnt* signal, highlighting the importance of the ventral *en1* domain and the capability of the appropriate *wnt* signal to arise dorsally. It is however important to consider that these dorsal limbs were derived from transplanted lateral plate mesoderm and not from a somitic cell population homologous to the median fins in which *wnt* signalling does not appear responsible for activation of *Lmx1b*.

**Fig. 5.**
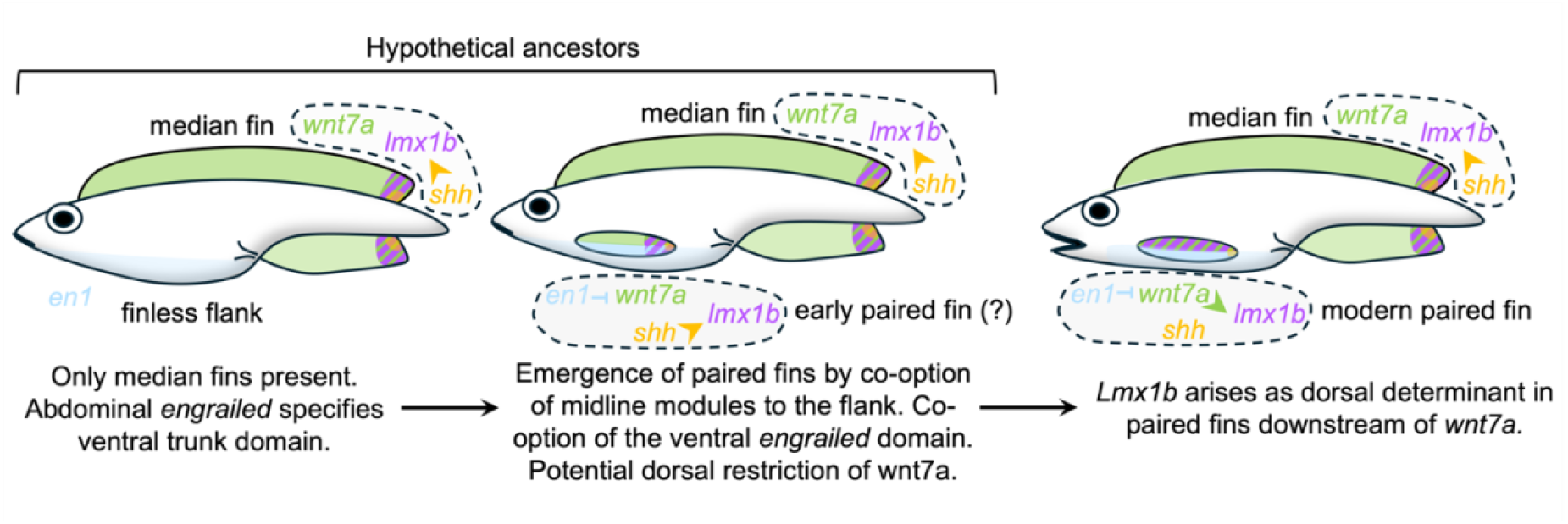
Model for the emergence of DV patterning in paired appendages. Ancestral median fins are predicted with posterior domain of Lmx1b downstream of shh from the ZPA as well as superficial wnt7a expression. Co-option of the median fins to the flank occurred on top of a pre-existing ventral domain of engrailed which became incorporated as the ventral side of the limb (Tanaka, et al. 2002; Tanaka and Onimaru 2012) and may directly have restricted wnt7a expression dorsally. Subsequent evolution of wnt responsive dorsal regulatory elements expanded Lmx1b expression in the dorsal side of the paired appendages.

In summary, we propose that *Lmx1b* was co-opted as dorsal determinant in paired fins and limbs from an original role in the posterior midline fins (Fig. 5). Although its exact function in this domain remains to be determined using targeted loss of function assays, we predict that activation of ephrin receptors for neuronal guidance is amongst them. Altogether, it appears that the DV axis of paired appendages should be interpreted as an evolutionary novelty that arose as amalgamation of ancestral midline determinants combined with novel regulatory inputs. This process provided the genetic substrate for the morphological individuation of dorsal and ventral limb surfaces and thereby enabled the evolution of terrestrial locomotion in land vertebrates.

## Material and Methods Embryo collection

Embryos of *A. burtoni* were collected from stocks at the TFA Uni-Konstanz and cultured as described (Höch, et al. 2021). The *gremlin1b* knockout mutant fish were generated as described using CRISPR/Cas9 under permit G18/32 (Höch, et al. 2021). The allele used in this study corresponds to a 86bp deletion within the open reading frame and is an alternate allele to the one reported before (Höch, et al. 2021). *A. baerii* embryos were obtained from a private breeder in Germany. Catharks were obtained from a zoo breeding station in Germany and as gift from Dr. Christina Paliou (Centro Andaluz de Biología del Desarrollo, Spain). All experimental procedures were performed and terminated before feeding stages rendering them exempt from further permission under German law.

### Cloning of Probes

Probes were cloned in pGEMT (Promega A3600) vector using a PCR template from synthesized embryonic cDNA of *A. burtoni, A. baerii* or *S. canicula*. Sequences of genes were identified by NCBI nucleotide BLAST against *A. burtoni, A. baerii* and *S. canicula* datasets. Primers were designed based on gene sequences deposited on NCBI. The primer table for each gene is provided in **Table S1**.

### *In situ* hybridization

*In situ* hybridization was performed according to (Woltering, et al. 2009; Woltering, et al. 2014; Woltering and Duboule 2015; Woltering, et al. 2020; Höch, et al. 2021) with the changes that material was stored in 100% Ethanol (to avoid Methanol toxicity) at -20°C after fixation, and the initial bleaching step was omitted. *A. baerii* 2 dph, 4dph and *A*.*burtoni* 8dpf embryos were bleached after completion of staining as follows: Embryos were washed extensively for a minimum of 2 days involving frequent buffer changes in TBST to remove all unconverted staining reagents (which would convert in contact with bleaching solution). Next, specimen were fixed with 4% PFA in PBS for 1-2 hours at room temperature to fix the converted stain, which otherwise diffuses in contact with organic solvents. Finally, pigmentation was removed using bleaching solution (1% H_2_O_2_, 5% Formamide, 0.5x SSC in H_2_O) by placing the samples under a bright white lamp for 1-2 hours. Afterwards specimen were washed with PBS and stored at 4°C in PBS supplemented with 0.02% NaN_3_ as preservative.

### Signal transduction modulation

Embryos were treated using the following concentrations of the different components: 1 μM SAG (Selleckchem S7779 - dissolved at 10 mM in DMSO), 5 μM cyclopamine (Selleckchem S1146 - dissolved at 50 mM in ethanol), 1 μM DMH1 (Selleckchem S7146 - dissolved at 20mM in DMSO), and 5 μM IWR-1-edo (Selleckchem S7086 - dissolved at 20 μM in DMSO). Embryos were cultured at a maximum density of 15 embryos per 100 ml dish with the usual supplementation of Penicillin-Streptomycin and methylene blue (Höch, et al. 2021). As the treatments were to be performed during the initial budding stages of the three different fin types, their heterochronic development had to be considered (Woltering, et al. 2018). For pectoral fins, embryos were treated from 2 – 4dpf, for pelvic fins, embryos were treated from 6 – 8dpf and for dorsal and anal fins, embryos were treated from 4 – 6 dpf (similar to the treatments performed in (Höch, et al. 2021)). All embryos were dechorionated before start of the treatments if necessary (applies to 2dpf/4dpf). Embryos were fixed in 4% PFA in PBS overnight (ON) at 4°C and stored in 100% Ethanol at -20°C.

### Paraffin embedding for histological sections

After staining, hybridized embryos were fixed overnight in PFA 4%, washed in PBS and dehydrated in increasing concentrations of ethanol, followed by two steps of xylene before being embedded in paraffin (SIGMA, #P-3558), where they were kept at 60°C overnight. The time for 100% EtOH and xylene steps were adjusted according to the stage and size of the sample. Histological sections of 7-8μm thickness were performed in paraffin-embedded samples using a Leica RM2125RT microtome and placed on SuperFrostPlus slides (LEICA, #10149870).

### Image acquisition

Images were acquired with a Leica MZ10F binocular microscope connected to a PC running LASv4.5 software. Images were assembled using the Z-stacking option. Colour images were acquired with a Leica DMC2900 camera; fluorescent images with a Leica DFC3000G camera and “ET UV LP” filter. For optimal contrast of non-stained embryo surfaces the *in situ* images shown in **Fig.2a, b**; **Fig. S2, Fig.S3, Fig.4b** left most panel “*ephA4b* posterior view” were constructed as overlays of a lower layer black-and-white fluorescent image of Hoechst stain (see below) and an upper transparent layer colour image (using the “layer opacity” function in Adobe Photoshop set between 70-90%). This method was previously used in (Woltering and Duboule 2015). All other images are presented as brightfield colour images with minimal adjustments to brightness and contrast. Images meant to show quantitative differences in expression (**Fig.3, Fig.4b, Fig.S5)** were taken with the same light and microscope settings and equally adjusted by (linear) brightness and contrast parameters in post between experimental and controls. For fluorescent images, embryos were stained after completion of the *in situ* in 2.5 μg/ml Hoechst stain (Sigma – B2262) in PBS with 0.02% NaN_3_ as preservative ON at 4°C. For consistent representation within the figure original images were sometimes reoriented and/or mirrored. The “stage indicating” *A*.*burtoni* embryos in **Fig.2a** were previously published in (Woltering, et al. 2018) under a Creative Commons Attribution 4.0 International License. Selective colour adjustments were made to the “stage indicating” *A. baerii* embryos in **Fig.2b** to eliminate background lighting artefacts that distorted the colour of the fin folds in the original images.

## Acknowledgements and funding

We thank David Walter for his technical assistance and Dr. Christina Palliou for the catshark specimens. JMW and SGB were supported by funds of the University of Konstanz and DFG project 404363160 (to JMW). SZN was supported by an EMBO Scientific Exchange Grant (#10496) for work carried out at the University of Konstanz. AM was supported by funds of the University of Konstanz. MR was supported by the Spanish Ministry of Science and Innovation (grant PID2023-147771NB-I00).

