## Supplementary Material for "Dorsoventral limb patterning in paired appendages emerged via regulatory repurposing of an ancestral posterior fin module"

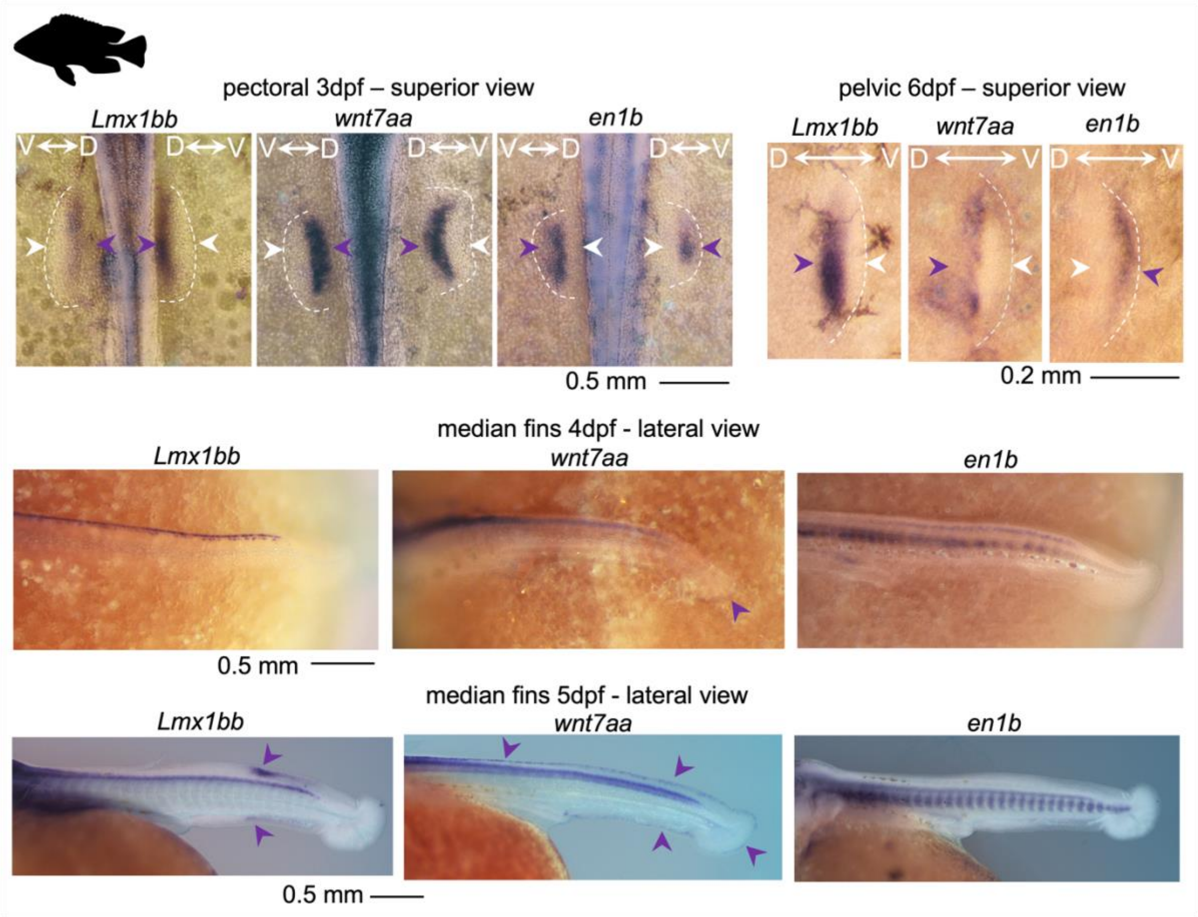


***Fig.S1) Expression of Lmx1bb, wnt7aa and en1b in A. burtoni 3-5 dpf embryos.***

*Similar dorsal restricted Lmx1bb/wnt7aa and ventrally restricted en1b expression can be observed at the earliest budding stages (3dpf/6dpf) of pectoral and pelvic fins. In 4dpf median fins only weak expression of wnt7aa can be detected in the emerging caudal fin. At 5dpf Lmx1bb has become activated in the dorsal and anal fins, whereby the dorsal fin appears slightly ahead in its development. At this stage wnt7aa is expressed superficially throughout the distal median fin Anlagen, forming a continuous domain including the fin folds (i.e. the parts between dorsal/anal and caudal fins that will not give rise to an adult fin). Lilac arrowheads indicate sites of notable expression while white arrowheads indicate absence of expression Abbreviations: dpf; days post fertilization*

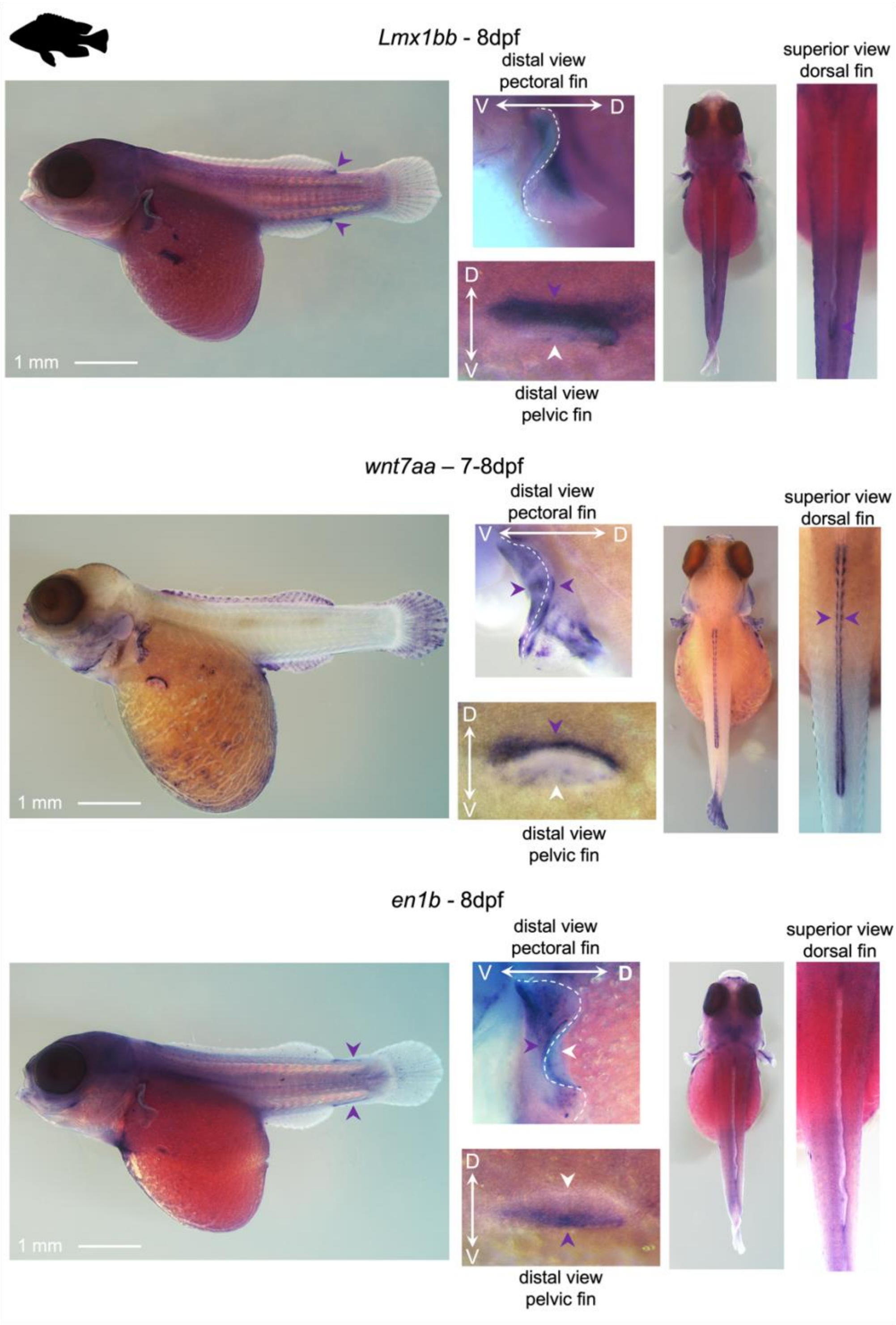


***Fig.S2) Expression of Lmx1bb, wnt7aa and en1b in A. burtoni 3-5 dpf embryos.***

*At 8dpf Lmx1bb expression is present as a medial proximal domain in the median and pectoral fins. Externally no clear DV polarity can be observed at this stage in pectoral fins. At this stage the Lmx1bb dorsal pattern is however obvious in the later emerging pelvic fins. Wnt7aa has become symmetrically expressed at this stage in pectoral fin rays, labelling both dorsal and ventral hemi-rays resembling the emerging lepidotrichia of the dorsal fin. This symmetric domain may be unrelated to its earlier superficial dorsal domain, at least the weak and diffuse expression of Lmx1bb at this stage in the distal fin suggest that wnt7aa is not capable of activating its expression at this stage. As for Lmx1bb, pelvic fins show clear dorsally restricted wnt7aa expression at this stage. En1b is not obviously expressed in any of the median fins but shows a surprising domain along the dorsal and ventral midline in between dorsal/anal and caudal fins. A superficial ventral domain appears preserved at this stage in pectoral fins. Pelvic fins clearly show ventrally restricted en1b expression at this stage. Lilac arrowheads indicate sites of notable expression while white arrowheads indicate absence of expression Abbreviations: dpf; days post fertilization.*

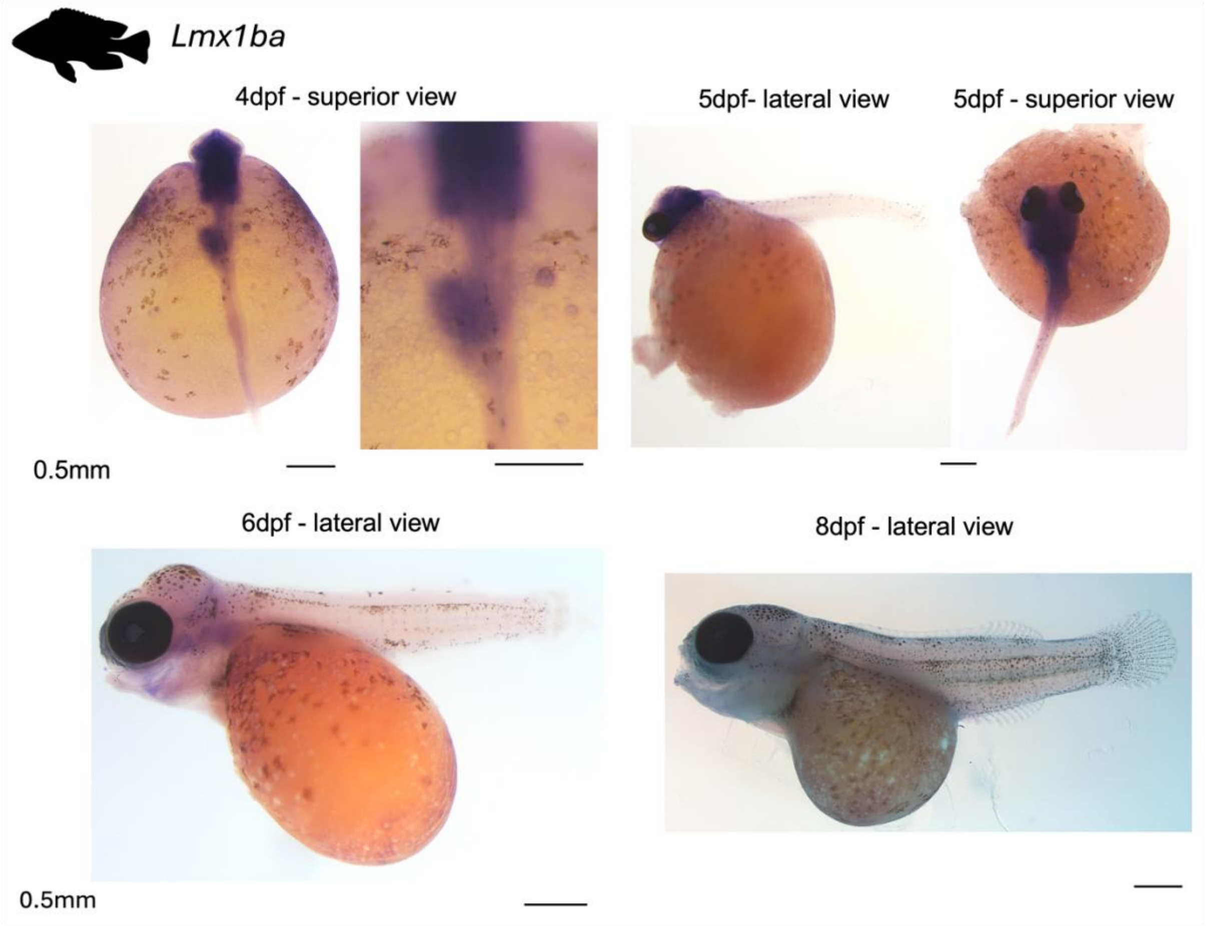


***Fig.S3) Expression of Lmx1ba in A. burtoni 4-8 dpf embryos.***

In contrast to *Lmx1bb*, *Lmx1ba* expression is not observed in fins at any stages. *Lmx1ba* is primarily expressed in neural tissue and in a domain in the forming gut at 4dpf (note overlap with *ephA4b* expression shown in **Fig.4).**

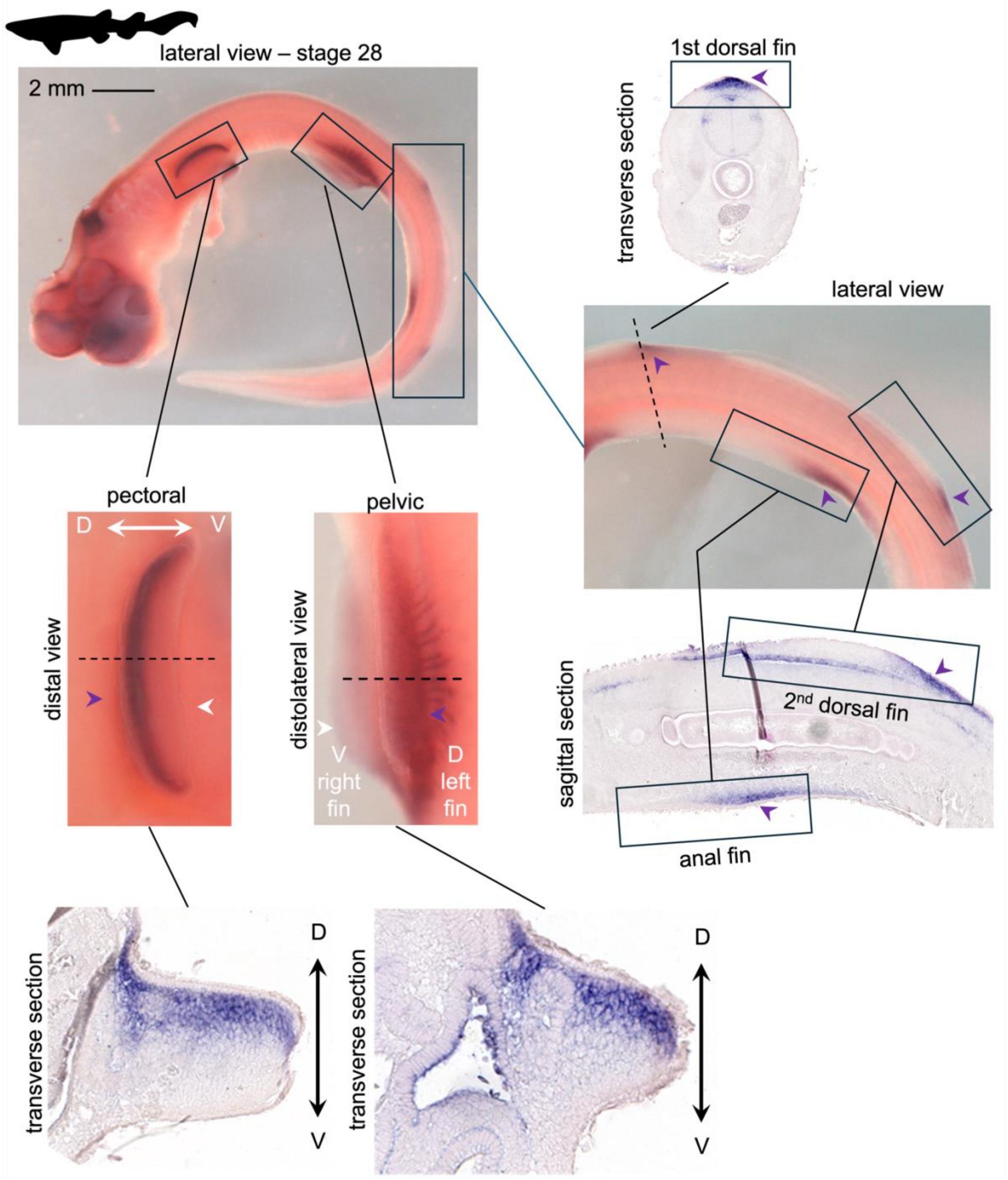


***Fig.S4) Expression of Lmx1b in S. canicula st.28.***

*Lmx1b is expressed with clear dorsal restriction in the pectoral and pelvic appendages of st.28 catshark as shown in whole mount pictures and transverse sections. In the median fins, 1st and 2nd dorsal and anal fins, Lmx1b expression initiates as a posterior median domain as shown by transverse section of the 1st dorsal fin and sagittal section of the 2nd dorsal and anal fin.*

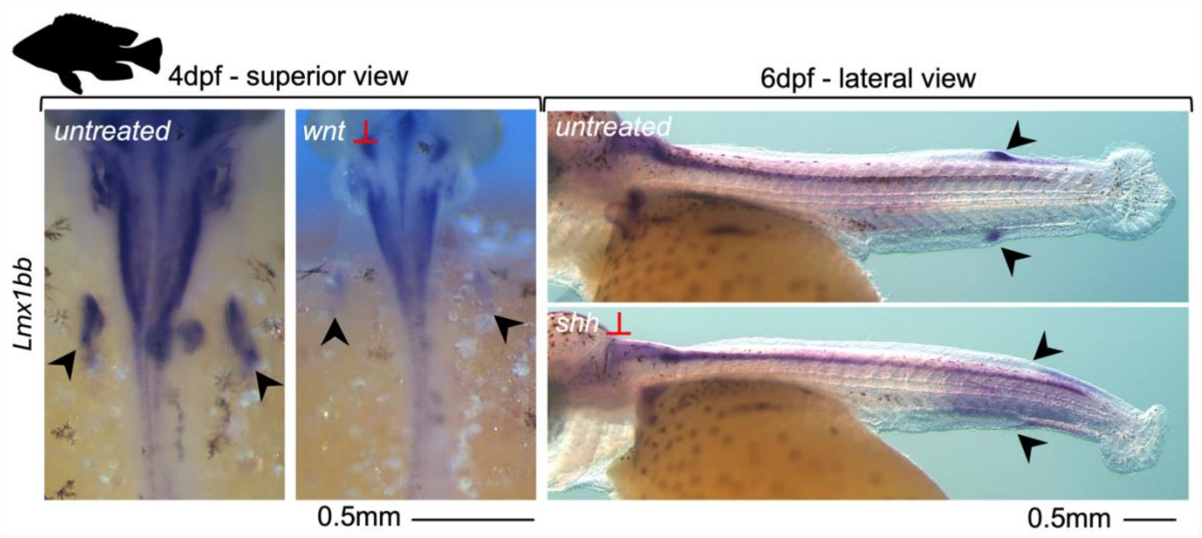


***Fig.S5) Downregulation of Lmx1bb in A.burtoni treated with wnt or shh inhibitors.***

*Embryos treated from 2-4 dpf with wnt signalling inhibitor show strong downregulation of Lmx1bb expression in pectoral fins as well as in neural structures. Embryos treated from 4-6 dpf with shh inhibitors show downregulation of the posterior Lmx1bb domain in dorsal and anal fins. The embryos shown function as control to the eph4Ab experiment in Fig.4b and were part of the same treatment. Dark arrowheads indicate pectoral fins in 4dpf and the presence/absence of posterior Lmx1bb expression at 6dpf.*

| Species | Gene | Primer Forward | Primer Reverse |
|---|---|---|---|
| ***A.burtoni*** | *Lmx1ba* | ACTACCAACAGTTGTTTGCTG | GGTAGGAGGTATGGTGGGCTG |
| | *Lmx1bb* | GACAAGCCACACTCGGAGTTATGC | GAGCCAGCGGTGTGAAGGAGTTC |
| | *Lhx2b* | CAGCATCTACTGCAAAGAGGAC | GCACTCGTGGACATCCATGCTAC |
| | *Lhx9* | GTCAGACCTGTGGAACACAG | TGCACCTCATCCAAGAACCAG |
| | *wnt7aa* | GTAAGGAGGCTGCTTTCACCTAC | CTATTTACAGGTATACACTTCTGTC |
| | *en1b* | ATGGAAGAGCAGAAGGAGCC | TCATTCGCTCTCCTCCTTCTCCTC |
| | *ephA4b* | GTTTACCTCAGCAAGTGATGTG | GTTGACACGGTGTGAAATGTC |
| ***A.baerii*** | *Lmx1b* | GAGAGCACTGGAGTGCGTCTATC | TCACGAAGCGAAGTACGAGTTCTG |
| | *Lhx2* | TGCTCGCTGTAGACAAACAGTG | GAAGAGACTCGTGAGGGTAG |
| | *Lhx9* | GTAGGGCAGAAGAAAACACTTG | GTGGTAGTGAAGTGCCATCAG |
| | *wnt7a* | GTTGCTCATGCCATCACAGCTG | CTGCAGGTGTTGCATTTGACGTAG |
| | *en1* | GCGCAGCAAGCTCACAGAAC | CCTGAACCGTGGTGGTGGAATG |
| ***S. canicula*** | *Lmx1b* | GTCCTGCCACCTTGGGAATG | CGTTCTGTTGTTCTTGCTGCTG |

**Table SI) Primers sequences used for probe cloning.**
